# Delivering on the Vision of Bench to Bedside: A Rare Disease Funding Community Collaboration to Develop Effective Therapies for Neurofibromatosis Type 1 Tumors

**DOI:** 10.1101/552976

**Authors:** Salvatore La Rosa, Vidya Browder, Annette C. Bakker, Jaishri O. Blakeley, Sharad K. Verma, Ling M. Wong, Jill A. Morris, Naba Bora

## Abstract

The time from target identification to new drug approval is often measured in decades. This can be even more challenging for rare diseases. Indeed, 95% of rare diseases do not have a specific therapy approved. Coordinated efforts to support research along the drug development pipeline can provide long term and comprehensive support to enable scientific breakthroughs for rare diseases. However, this requires coordination across multiple stakeholders. The present article analyzes the funding efforts of four major federal and philanthropic organizations to accelerate the advancement of MEK inhibitors to human clinical trials for NF1-associated tumors.

## Introduction

Neurofibromatosis type 1 (NF1) is an autosomal dominant genetic syndrome caused by mutations in the *NF1* gene (Gutmann, et al. 2017). Development of multiple tumors, including plexiform neurofibromas (pNF) and gliomas, are hallmarks of NF1. There are multiple additional manifestations and the clinical severity of NF1 is highly variable, with significant differences even within families sharing a mutation. NF1 patients are also at increased risk of developing malignancies such as malignant peripheral nerve sheath tumors (MPNST) and breast cancer (Frayling, et al. 2018). Despite their significant morbidity, therapeutic development for NF1 has been slow and challenging because of an incomplete understanding of the disease biology. Although surgical debulking is the preferred treatment option for pNFs, it can be extremely complicated, especially if the tumor is intricately involved with nerve tissue, vasculature, or a vital organ. Surgery is rarely performed for NF1 optic pathway gliomas because of their location. Symptomatic gliomas are typically managed with chemotherapeutic agents, which, though effective, are associated with long term sequelae such as cognitive dysfunction (de Blank, Berman and Fisher 2016). Thus, an effective long-term therapy for NF1 associated tumors is an unmet medical need.

The *NF1* protein product, neurofibromin, negatively regulates Ras signal transduction pathways. Several preclinical studies validated mitogen-activated protein kinase (MEK), a component of the Ras-Raf-MEK-ERK signaling cascade, as a potential therapeutic target for NF1 tumors and other manifestations. At least four MEK inhibitors, PD0325901, trametinib (GSK1120212, Mekinist), binimetinib (ARRY-438162, MEK162), and selumetinib (AZD6244), have progressed to clinical trials in NF1 patients and, at the time of writing, ClinicalTrials.gov lists 19 trials where these MEK inhibitors are being evaluated for various manifestations of NF1 (Table 1). In fact, selumetinib was granted Orphan Drug Designation by the US Food and Drug Administration (FDA) and the European Medicines Agency (EMA) based on encouraging results in pediatric NF1 patients with inoperable pNFs (Dombi, et al. 2016). If approved, selumetinib will be the first drug available for the treatment of NF1-associated tumors.

**TABLE 1.** Ongoing MEK inhibitor clinical trials for conditions including NF1*.

| **ClinicalTrials.gov identifier** | **Intervention** | **Study name** | **Phase** | **Recruitment status** |
| --- | --- | --- | --- | --- |
| NCT02096471 | PD0325901 | MEK Inhibitor PD-0325901 Trial in Adolescents and Adults With NF1 (MEK Inhibitor) | Phase 2 | Active, not recruiting |
| NCT02124772 | Trametinib, Dabrafenib | Study to Investigate Safety, Pharmacokinetic (PK), Pharmacodynamic (PD) and Clinical Activity of Trametinib in Subjects With Cancer or Plexiform Neurofibromas and Trametinib in Combination With Dabrafenib in Subjects With Cancers Harboring V600 Mutations | Phase 1/2a | Recruiting |
| NCT03363217 | Trametinib | Trametinib for Pediatric Neuro-oncology Patients With Refractory Tumor and Activation of the MAPK/ERK Pathway | Phase 1/2 | Not yet recruiting |
| NCT03190915 | Trametinib | Trametinib in Treating Patients With Relapsed or Refractory Juvenile Myelomonocytic Leukemia | Phase 2 | Recruiting |
| NCT03232892 | Trametinib | Trametinib in Patients With Advanced Neurofibromatosis Type 1 (NF1)-Mutant Non-small Cell Lung Cancer | Phase 2 | Recruiting |
| NCT02465060 | Trametinib | Targeted Therapy Directed by Genetic Testing in Treating Patients With Advanced Refractory Solid Tumors, Lymphomas, or Multiple Myeloma (The MATCH Screening Trial) | Phase 2 | Recruiting |
| NCT01885195 | Binimetinib | MEK162 for Patients With RAS/RAF/MEK Activated Tumors (SIGNATURE) | Phase 2 | Recruitment completed |
| NCT03231306 | Binimetinib | Phase II Study of Binimetinib in Children and Adults With NF1 Plexiform Neurofibromas (NF108-BINI) | Phase 2 | Recruiting |
| NCT02285439 | Binimetinib | Phase I/II Study of MEK162 for Children With Ras/Raf Pathway Activated Tumors | Phase 1/2 | Recruiting |
| NCT01089101 | Selumetinib | Selumetinib in Treating Young Patients With Recurrent or Refractory Low Grade Glioma | Phase 1/2 | Recruiting |
| NCT01362803 | Selumetinib | AZD6244 Hydrogen Sulfate (Selumetinib) for Children With Nervous System Tumors | Phase 2 | Recruiting |
| NCT02407405 | Selumetinib | Selumetinib in Treating Patients With Neurofibromatosis Type 1 and Plexiform Neurofibromas That Cannot Be Removed by Surgery | Phase 2 | Recruiting |
| NCT02839720 | Selumetinib | Selumetinib in Treating Patients With Neurofibromatosis Type 1 and Cutaneous Neurofibroma | Phase 2 | Recruiting |
| NCT03109301 | Selumetinib | Mitogen Activated Protein Kinase Kinase (MEK1/2) Inhibitor Selumetinib (AZD6244 Hydrogen Sulfate) in People With Neurofibromatosis Type 1 (NF1) Mutated Gastrointestinal Stromal Tumors (GIST) | Phase 2 | Recruiting |
| NCT03213691 | Selumetinib | Selumetinib in Treating Patients With Relapsed or Refractory Advanced Solid Tumors, Non-Hodgkin Lymphoma, or Histiocytic Disorders With Activating MAPK Pathway Mutations (A Pediatric MATCH Treatment Trial) | Phase 2 | Recruiting |
| NCT03259633 | Selumetinib | An Intermediate Access Protocol for Selumetinib for Treatment of Neurofibromatosis Type 1 | Expanded access | Available |
| NCT03326388 | Selumetinib | Intermittent Dosing Of Selumetinib In Childhood NF1 Associated Tumours (INSPECT) | Phase 1/2 | Not yet recruiting |
| NCT03649165 | Selumetinib | A Study to Evaluate Bioavailability and Food Effect of Selumetinib (AZD6244) in Healthy Male Participants | Phase 1 | Not yet recruiting |
| NCT03433183 | Selumetinib | SARC031: MEK Inhibitor Selumetinib (AZD6244) in Combination With the mTOR Inhibitor Sirolimus for Patients With Malignant Peripheral Nerve Sheath Tumors | Phase 2 | Not yet recruiting |

\*Search done on clinicaltrial.gov on Feb 1, 2019.

The funding landscape of NF1-MEK research, and of NF1 research as a whole, has been shaped by the efforts of multiple federal and philanthropic organizations. In addition to the National Institutes of Health (NIH, intramural and extramural), the organizations and research programs that have played a crucial role in advancing NF1-MEK research include the Department of Defense (DoD) Congressionally Directed Medical Research Programs (CDMRP), Children’s Tumor Foundation (CTF), and Neurofibromatosis Therapeutic Acceleration Program (NTAP) at Johns Hopkins University. This report presents a retrospective analysis of the NF1-MEK funding data for the years 2006-2017 including, but not limited to, funding from the above mentioned organizations. Through this analysis, the authors illustrate how the contributions across the funders during the past 10 years have made the availability of a drug for NF1 tumors a plausible reality.

## Methods

To extract all available NF1-MEK funding data, Dimensions for Funders (https://www.dimensions.ai/), a database of publicly and privately funded research projects worldwide, was used. The database was queried using an intersect of two queries that were each specific for MEK inhibitor-related and NF1-related grants. The search was restricted to 2006-2017 to ensure completeness of data among the relevant funders. Results from this search were supplemented using iSearch, a portfolio analysis tool internal to NIH, and Research, Condition, and Disease Categorization (RCDC), a classification scheme used by the NIH for reporting (NIH, 1998) (NIH, 2011). The RCDC classification scheme has had a neurofibromatosis category that includes NF1, NF2, and schwannomatosis since 2008. As a result, there was no neurofibromatosis category data from 2006-2008 available for this analysis. iSearch was used to query the neurofibromatosis RCDC category. The integrated output across datasets was manually inspected for relevance to MEK inhibitor development to treat NF1, yielding a final curated list of NF1-MEK grants. The Broad Research Area (BRA) classification implemented in Dimensions was used to distribute the grants into two categories: ‘basic research’ and ‘clinical research’. **Basic Research**: This classification includes research grants from pure basic science to applied research including *in-vitro* studies. **Clinical Research**: This classification includes research grants from *in-vivo* exploratory studies through clinical trials. Several analysis details are notable: 1) Almost all grants are multi-year funding, and the total amount reported here includes committed future costs for ongoing grants; 2) Foreign grants were converted to USD based on the exchange rate at the time of the award; 3) Multi-year projects that began before 2006 were not included, even if they were active in 2006. (TABLE 2)

**TABLE 2.** Federal and philanthropic research organizations providing NF1-MEK research funding in 2006-2017.

| FUNDER | GRANTS IN 2006-2017 |  | BASIC RESEARCH |  | CLINICAL RESEARCH |  |
| --- | --- | --- | --- | --- | --- | --- |
|  | Number | Funding (USD) | Number | Funding (USD) | Number | Funding (USD) |
| NIH | 29 | 51,815,350 | 17 | 25,114,554 | 12 | 26,700,796 |
| CDMRP | 17 | 38,448,301 | 7 | 4,004,176 | 10 | 31,444,125 |
| CTF | 34 | 6,020,624 | 20 | 1,374,947 | 14 | 4,645,677 |
| NTAP | 5 | 3,755,487 | - | - | 5 | 3,755,487 |
| CTF/NTAP (co-funding) | 1 | 3,658,649 | - | - | 1 | 3,658,649 |
| Others | 13 | 1,723,644 | 8 | 641,486 | 5 | 1,082,158 |
| Total | 99 | 102,422,055 | 52 | 31,135,163 | 47 | 71,286,892 |

## Results

We identified a total of 99 NF1-MEK grants for 2006-2017 with an aggregate funding of $102.42 million. Of these, 86 grants totaling $100.7 million were funded by the NIH, CDMRP, CTF, and NTAP, making these four organizations the largest funders in this area. The remaining 13 grants for a total of $1.7 million were awarded by four organizations: the Japan Society for the Promotion of Science, Melanoma Research Alliance, St. Baldrick’s Foundation, and National Natural Science Foundation of China.^1^

**FIGURE 1.**
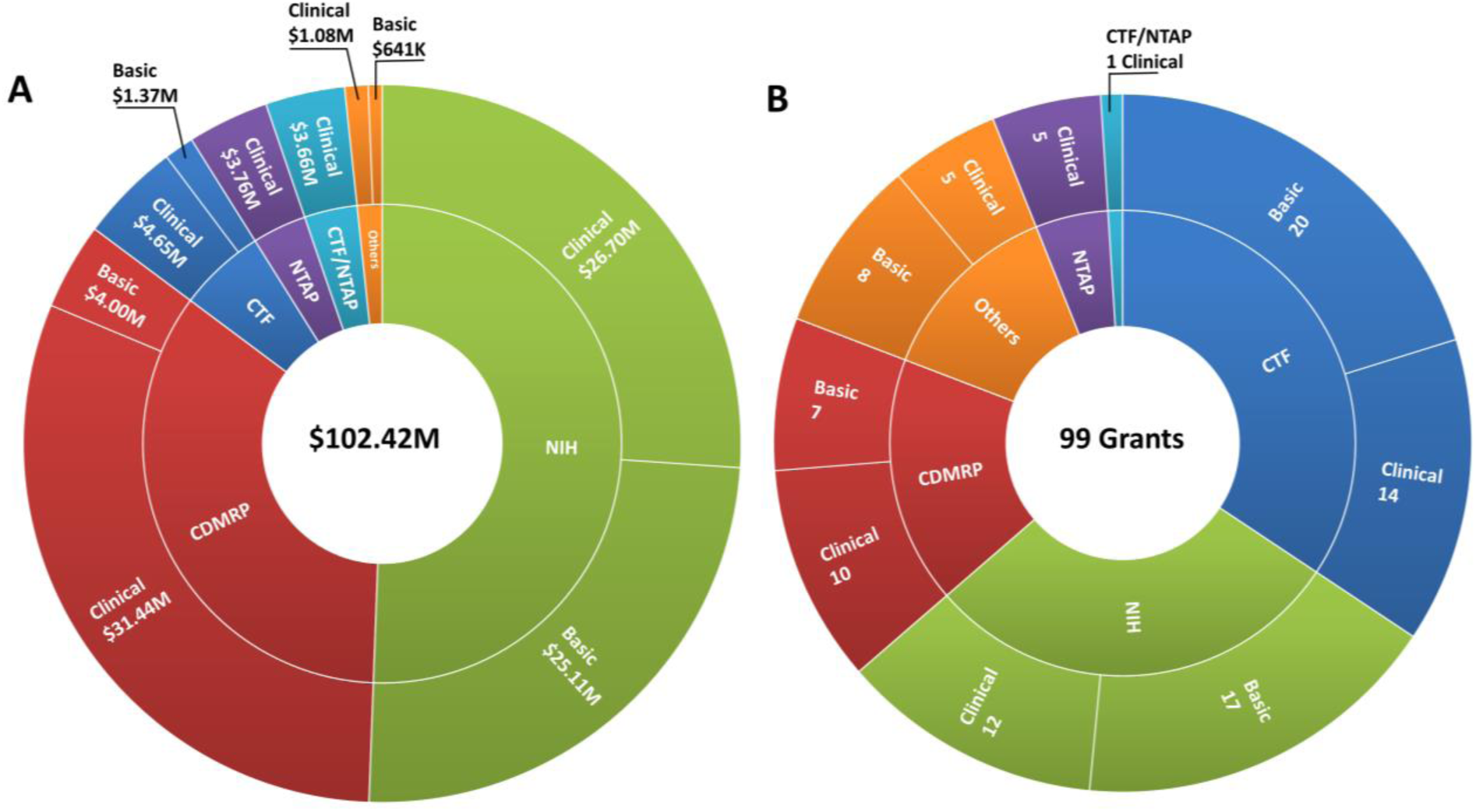
Total funding across organizations for NF1-MEK research in 2006-2017 A) Funding amounts. **B)** Number of grants.

### National Institutes of Health

The NIH was the leading federal funder of NF1-MEK research, with $51.8 million granted through 29 grants. NIH grants included eleven R01 grants totaling $18.6 million, four National Cancer Institute (NCI) intramural research projects involving clinical trials or natural history studies totaling $18.4 million, a bench-to-bedside award, a Javits award, and several training grants. Funding was approximately evenly distributed between basic and clinical research ($25.1 vs $26.7 million, respectively). Relative to the other NIH institutes and centers, the NCI and the National Institute of Neurological Disorders and Stroke (NINDS) have been most involved in the development of MEK inhibitors and as applied to NF1. Specifically, in 2006-2017 the NCI awarded $33.9 million between its intramural and extramural programs, the NINDS awarded $15.4 million, and remaining NIH funding was provided by the National Institute on Deafness and Other Communication Disorders, National Institute of Arthritis and Musculoskeletal and Skin Diseases (NIAMS), and National Institute of Dental and Craniofacial Research (NIDCR). In addition to funding several basic research studies in NF1, the NCI has conducted or funded many of the MEK clinical trials, including the first clinical trial to evaluate a MEK inhibitor (CI-1040; NCT00033384 and NCT00034827), the first selumetinib trial (NCT01089101), five other selumetinib trials (NCT01362803, NCT02407405, NCT02839720, NCT03109301, NCT03213691), two other interventional trials (NCT03190915, NCT02465060), and an ongoing natural history study. The NINDS has primarily supported preclinical studies and a national, interdisciplinary center for NF research. NINDS-funded pre-clinical contributions include establishment of a murine model of NF1-related MPNSTs, development of preclinical screening tests to compare potential therapeutics in NF1 cell lines, and completion of proof-of-efficacy studies for PD0325901.

### Neurofibromatosis Research Program (NFRP)

The NFRP at CDMRP was identified as the second largest federal contributor to NF1-MEK funding, having awarded approximately $35.4 million across 17 grants in 2006-2017. Nearly 89% of the dollar amount was directed towards clinical studies. Unlike the NIH, the NFRP allocates a congressionally appropriated budget each year for all forms of NF research. Through the NF Clinical Trials Consortium (NFCTC), the CDMRP has invested more than $27 million in developing all the infrastructure of the consortium and conducting 10 interventional trials, including two trials with MEK inhibitors in pNFs (PD0325901, NCT02096471) and Low-Grade Gliomas (binimetinib, NCT02285439). In addition to initiating clinical trials, the NFRP also funded eight Investigator-Initiated Research Awards ($7.2 million) and five Exploration – Hypothesis Development Awards ($726,504). Other funding mechanisms include the New Investigator Award to encourage non-NF investigators to conduct research in this area, and the Clinical Trial Award (1 award, $540,104).

### Children’s Tumor Foundation

The CTF was the leading non-federal philanthropic funder of NF1-MEK research in 2006-2017, during which time it awarded almost $6.2 million through 34 grants. As the oldest US-based philanthropic foundation dedicated to all forms of NF, CTF has historically acted as a discovery research seeder and gap funder through its funding mechanisms such as the Young Investigator Award, Drug Discovery Award, and Clinical Research Award. Of the 34 grants, 17 ($1.4 million) were Young Investigator Awards (YIA), ten ($371,123) were Drug Discovery Initiatives (DDI), five ($664,000) were Clinical Research Awards (CRA) and one ($159,500) was a Contract Award (CA). The CTF’s Neurofibromatosis Preclinical Consortium (NFPC, funded from 2008-2013) awarded almost $3.4 million for preclinical studies, followed by the Neurofibromatosis Therapeutic Consortium (NFTC, funded conjointly with NTAP from 2013-2016) that continued funding for $3.6 million. During the 2008-2013 period, the two consortia conducted more than 95 preclinical studies of 38 drugs or drug combinations through collaborations with 18 pharmaceutical companies (Maertens, et al. 2017). Over 40 studies included a MEK inhibitor either as a single agent or in combination (unpublished results). These programs generated critical data in support of MEK as a valid target for *NF1*-driven tumors, providing the preclinical basis that justified the initiation of clinical trials (NCT02096471, NCT02124772, NCT02285439, NCT01362803, NCT02407405) with MEK inhibitors (Chang, et al. 2013) (Jessen, et al. 2013).

### Neurofibromatosis Therapeutic Acceleration Program

The NTAP, although founded only in 2012, has emerged as the second largest non-federal non-profit funder of NF1-MEK research. Focused specifically on accelerating the development of therapeutics for pNF and cutaneous neurofibromas (cNF), since its inception NTAP has teamed up with CTF for the co-sponsoring of preclinical testing through the NFTC consortium. It has also invested over $3.7 million in MEK research, playing a critical role in facilitating the clinical evaluation of selumetinib in children with pNF (NCT02407405).

**FIGURE 2:**
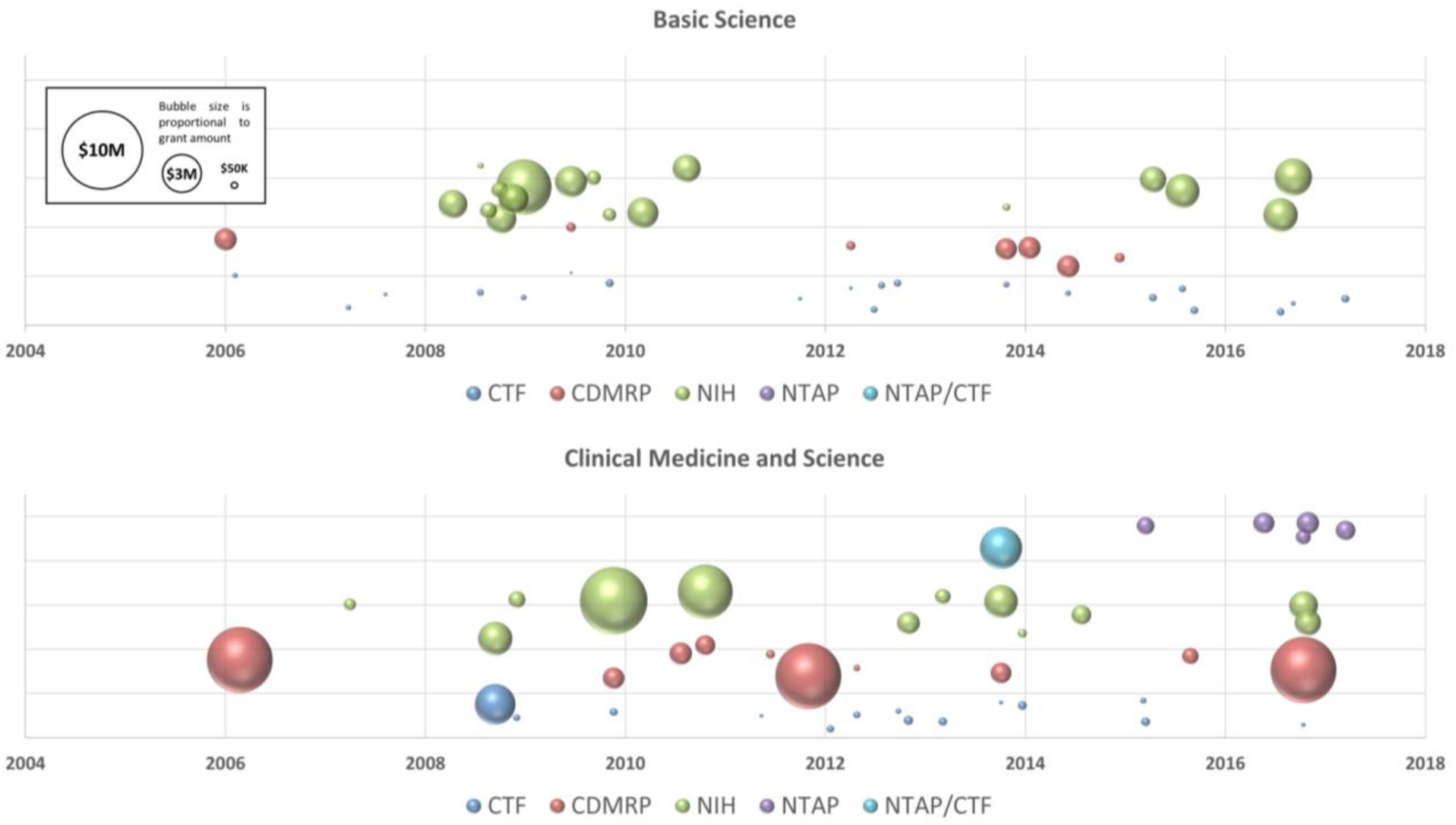
Timeline of NF1-MEK research awards by major funding organizations. Legend: Bubble size is proportional to the grant size and each grant is placed on the timeline based on its award date. The position of bubbles is jittered on the graph to facilitate visualization. Almost all grants are multi-year funding; for display purposes, the total amount is attributed to the initial year.

## Discussion

The retrospective analysis of the MEK funding landscape of the past 10 years has shown that the critical NF funders (NIH, CDMRP, CTF, and NTAP) have played distinct and complementary roles in the success of developing a MEK inhibitor to treat NF1 associated tumors. While NIH and CDMRP have provided the bulk of research dollars across all stages of NF research, CTF has seed funded NF research with a stream of small grants in the basic and translational research area. At the same time, funding for over $7 million in preclinical studies (NFPC and NFTC) allowed researchers to gather essential information to start clinical studies. Specific funders, like NIH and NTAP, have catalyzed critical steps and funded or co-funded preclinical or clinical stage projects that were essential to both launch and complete MEK clinical trials in people with NF1.

The NF1-MEK example illustrates the net outcome of the highly complementary approaches of the funding agencies and the yield of sustained funding investments across all stages of research. Overall, these programs and grants have enabled a MEK inhibitor to move along the drug discovery pipeline through clinical trials and to hopefully demonstrate its safety and effectiveness for patients in the near future.

The Dimension for Funders platform and similar platforms allow transparency in funding that allow stakeholders to coordinate efforts. This is particularly valuable in a rare disease community to maximize resources and ensure that follow on studies that are necessary to realize clinical benefit are pursued. Such integrated funding data also allows funders to identify gaps and overlaps in their investments so they work more effectively together to foster a community of researchers that has the tools and knowledge to advance NF research. As a next step, all NF funders are now collaborating on the largest data sharing effort ever initiated for NF. Spearheaded by CTF in 2014, all NF funders are now incentivizing their applicants to share their NF data openly on the recently launched NF data portal (www.nfdataportal.org). The portal is part of the largest NF Open Science Initiative (NF-OSI) to support the NF data discovery needs of computational biologists, bench scientists, clinicians, and the public. Participants in the NF-OSI benefit from early access to data and support for data sharing and re-use within the NF community.

Understanding the unique potential of each funding organization and leveraging each other’s strengths will bring us close to the next big discovery.

## Acknowledgments

The authors wish to acknowledge the following staff for their contributions to this analysis: Kimberly Pope and Whitney Cabrera from the NFRP/CDMRP, Cara Long from the Office of Science Policy and Planning at NINDS.

## Footnotes

1 Search executed on Oct 30, 2018

